# Laminar-specific functional connectivity mapping with multi-slice line-scanning fMRI

**DOI:** 10.1101/2021.03.03.433376

**Authors:** Sangcheon Choi, Hang Zeng, Yi Chen, Filip Sobczak, Chunqi Qian, Xin Yu

## Abstract

Laminar BOLD-fMRI has been applied to better depict the neuronal input and output circuitry and functional connectivity across cortical layers by measuring local hemodynamic changes. Despite extensive studies detecting laminar fMRI signals to illustrate the canonical microcircuit, the spatiotemporal characteristics of laminar-specific information flow across different cortical regions remain to be fully investigated in both evoked and resting states. Here, we developed a multi-slice line-scanning fMRI (MS-LS) method to detect laminar fMRI signals in adjacent cortical regions with high spatial (50 µm) and temporal resolution (100 ms) in anesthetized rats. Across different scanning trials, we detected both laminar-specific positive and negative BOLD responses in the surrounding cortical region adjacent to the most activated cortex under evoked condition. Specifically, in contrast to the typical Layer (L) 4 correlation across different regions due to the thalamocortical projections for trials with positive BOLD, a strong correlation pattern specific in L2/3 was detected for the trials with negative BOLD in adjacent regions, which indicate a brain state-dependent laminar-fMRI responses based on cortiocotical interaction from different trials. Also, we acquired the laminar-specific rs-fMRI signals across different cortical regions, of which the high spatiotemporal resolution allows us to estimate lag times based on the maximal cross-correlation of laminar-specific rs-fMRI signals. In contrast to the larger variability of lag times in L1 and 6, robust lag time differences in L2/3, 4, and 5 across multiple cortices represented the low-frequency rs-fMRI signal propagation from the caudal to the rostral slice. In summary, our work provides a unique laminar fMRI mapping scheme to better characterize trial-specific intra- and inter-laminar functional connectivity with MS-LS, presenting layer-specific spatiotemporal variation of both evoked and rs-fMRI signals.

## INTRODUCTION

Resting-state (rs-) fMRI is widely utilized in clinical and cognitive neuroscience for functional connectivity mapping, attributing to its unique capability to identify low frequency oscillation arised from neuronal oscillation across different brain states (1-5). The conventional method of rs-fMRI is based on echo-planar-imaging (EPI) to detect T2*-weighted blood-oxygen-level-dependent (BOLD) signal low-frequency fluctuation across functionally connected brain regions (6). This mapping method has been increasingly utilized to parcellate functional networks across the whole brain (7-11). Lately, high field fMRI has also been utilized in the mapping scheme to identify dynamic BOLD responses and cerebral blood volume (CBV) signals with layer specificity in bottom-up or top-down tasks (12-18), indicating the possible functional network across different cortical layers. However, the spatial resolution of fMRI images is often limited by the demanding requirement for high temporal resolution, i.e., fast-sampling of rs-fMRI oscillation patterns, at sub-second with sufficient signal-to-noise ratio (SNR) (19-21). In addition, potential aliasing effects from the cardiorespiratory artifacts (22-24) contaminate fMRI signals, leading to spurious results for rs-fMRI studies without sufficient sampling rates. It remains challenging to achieve high spatial resolution with a fast sampling rate to acquire rs-fMRI signals with sufficient SNR for brain functional mapping.

Previously, Yu et al. have developed a line-scanning fMRI method to substantially improve spatiotemporal resolution in rat brains, achieving 50-µm spatial resolution along a line profile which covers different cortical layers within 50-ms TR (25). Line-scanning fMRI has also been used in combination with fiber-based optogenetic stimulation (26) or diffusion-sensitizing gradients (27) to study the fast functional onset with high temporal resolution. Beyond preclinical studies, line-scanning fMRI is also being utilized in human fMRI studies to provide functional maps of cortical layers in a reduced field-of-view (FOV) with high spatiotemporal resolution (28-30). Lately, a similar line-scanning strategy has been used to map diffusion signals in the human brain (31). To date, however, no fMRI measurements have been performed on multiple cortical regions manifesting laminar-specific functional connectivity with high spatiotemporal resolution comparable to the original line-scaninng fMRI method.

In this work, we extended the line-scanning fMRI method towards multi-slice acquisition to acquire line profiles from different cortical regions with high spatial (50 µm) and temporal (100 ms) resolution. To investigate functional connectivity driven by the underlying neural network, we analyzed the intra- and inter-laminar correlation features under both evoked and resting-state conditions in anesthetized rats. Upon electrical stimulation of the forepaw, adjacent cortical regions to the forepaw somatosensory cortex (FP-S1) showed either positive or negative BOLD responses with distinct laminar-specific correlation patterns across different fMRI trials. Meanwhile, owing to the fast sampling rate, we were able to detect laminar-specific lag times of rs-fMRI signals with low-frequency fluctuation, showing specific signal propagation (< 0.1 Hz) across different cortices. Our work demonstrates the feasibility of multi-slice line-scanning fMRI (MS-LS) to identify evoked laminar correlation patterns and resting-state signal propagation across different cortical regions.

## RESULTS

### Mapping evoked laminar-specific BOLD signals with multi-slice line-scanning fMRI

We first developed the MS-LS method (**Fig. 1A**) to map laminar-specific BOLD responses from different cortical regions covering FP-S1 and adjacent cortices in one hemisphere (25) (details in **Method**). Left forepaw stimulation (**Fig. 1B**) activated contralateral cortical FP-S1, which has been reliably identified by using the conventional echo-planar-imaging (EPI) method (**Fig. 1A**) (32-34). As shown in **Fig. 1C**, three different line-scanning profiles from the corresponding cortical regions (i.e., rostral, middle, and caudal slices) were acquired by switching off phase-encoding gradient and defining the field-of-view (FOV) with two saturation slices to avoid aliasing from regions outside the FOV (**Fig. 1A**). Evoked BOLD-fMRI signals in three slices were simultaneously recorded as a function of time across the cortical depth (0-2 mm), showing the most salient BOLD responses in the caudal slice with decreased BOLD signals from the middle to the rostral slice as shown in **Fig. 1D-F**.

**Figure 1.**
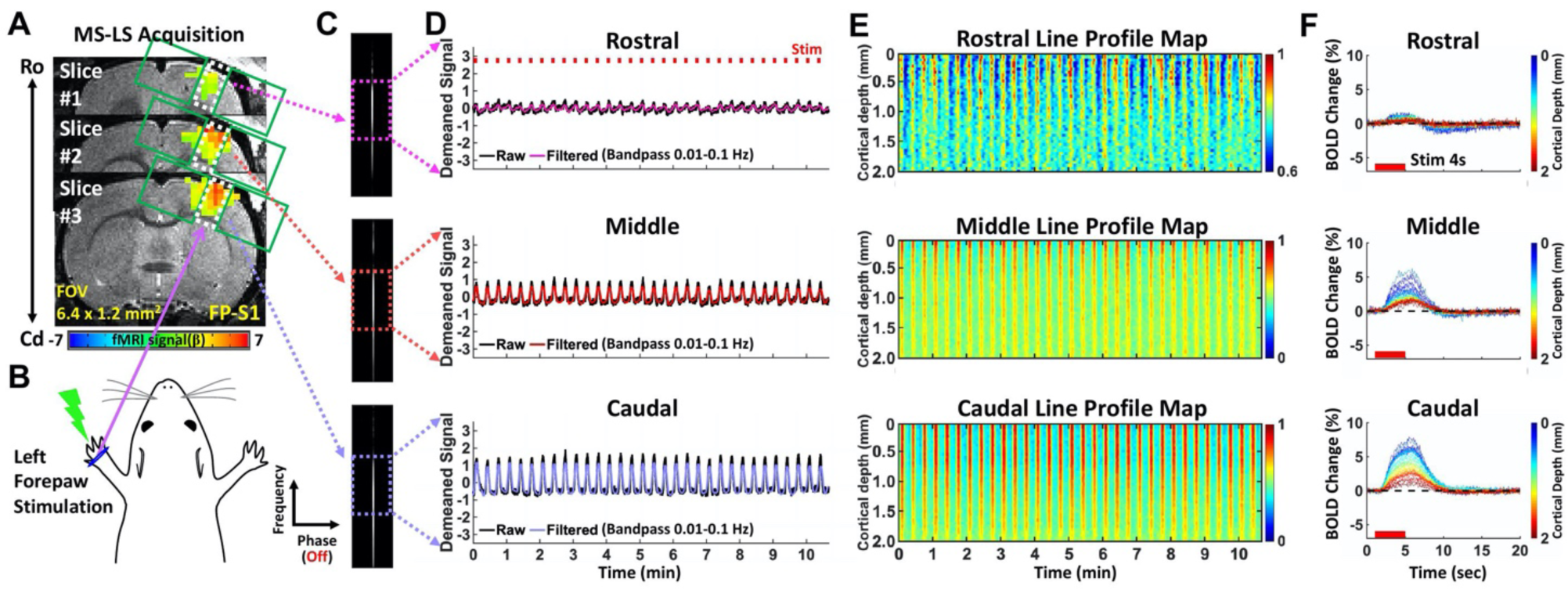
Evoked BOLD responses using the MS-LS method. **A**. Schematic illustration of the multi-laminar line-scanning experimental design on EPI-BOLD activation maps of different cortical regions overlaid on anatomical RARE images. Three different regions (white boxes) between two saturation slices (green boxes) were targeted from a rostral (Ro) to a caudal (Cd) region covering FP-S1 of one hemisphere. **B**. Left forepaw electrical stimulation (3 Hz, 4 s, 2.5 mA) following an fMRI design paradigm (1 s off, 4 s on and 15 s off). **C**. Representative three different line-scanning profiles from the corresponding slices. Each colored box indicates the corresponding cortical region. **D-F**. Average results from all data sets (n = 18 trials of 4 rats). **D**. Demeaned fMRI time series of raw (black) and filtered (magenta, orange, light purple, bandpass: 0.01-0.1 Hz) data (average of 40 voxels) in the rostral (upper), middle (middle), and caudal (lower) cortical regions under the evoked condition. Red boxes indicate 32 epochs for 10 min 40 sec. **E**. Normalized line profile maps showing the laminar-specific fMRI responses across the cortical depth (0–2 mm) in the three corital regions (40 voxels with 50 µm resolution). The time courses were processed by band-pass filtering (0.01-0.1 Hz). **F**. Average BOLD changes of the individual voxel time courses (mean epoch with 20 s) across the cortical depth in the three cortical regions.

### Distinguishing the inter- and intra-laminar functional connectivity across different cortices

Besides the typical positive BOLD signal detected in FP-S1 and adjacent area upon stimulation (25,26,35), negative BOLD signals were also reported in the adjacent cortex of activated regions (36-40). We applied MS-LS to characterize the laminar-specific correlation patterns of varied BOLD signals in cortical areas adjacent to activated FP-S1. As shown in **Fig. 2A**, highly varied BOLD responses in the rostral slice were observed across all trials with left forepaw stimulation. We sorted individual trials into two groups based on BOLD responses of the rostral slice, showing group 1 with positive and group 2 with negative BOLD signals (**Fig. 2A** and S1). **Fig. 2B** shows the laminar-specific BOLD responses across the three different slices for both groups. The spreading negative BOLD responses were observed at the superficial layers in the rostral slice and salient undershoot of the BOLD signal was detected in the middle slice in group 2 in contrast to the monophasic positive BOLD responses across the three slices in group 1. This result demonstrates that distinct BOLD responses in adjacent cortices to activated FP-S1 can be specified by MS-LS with laminar specificity.

**Figure 2.**
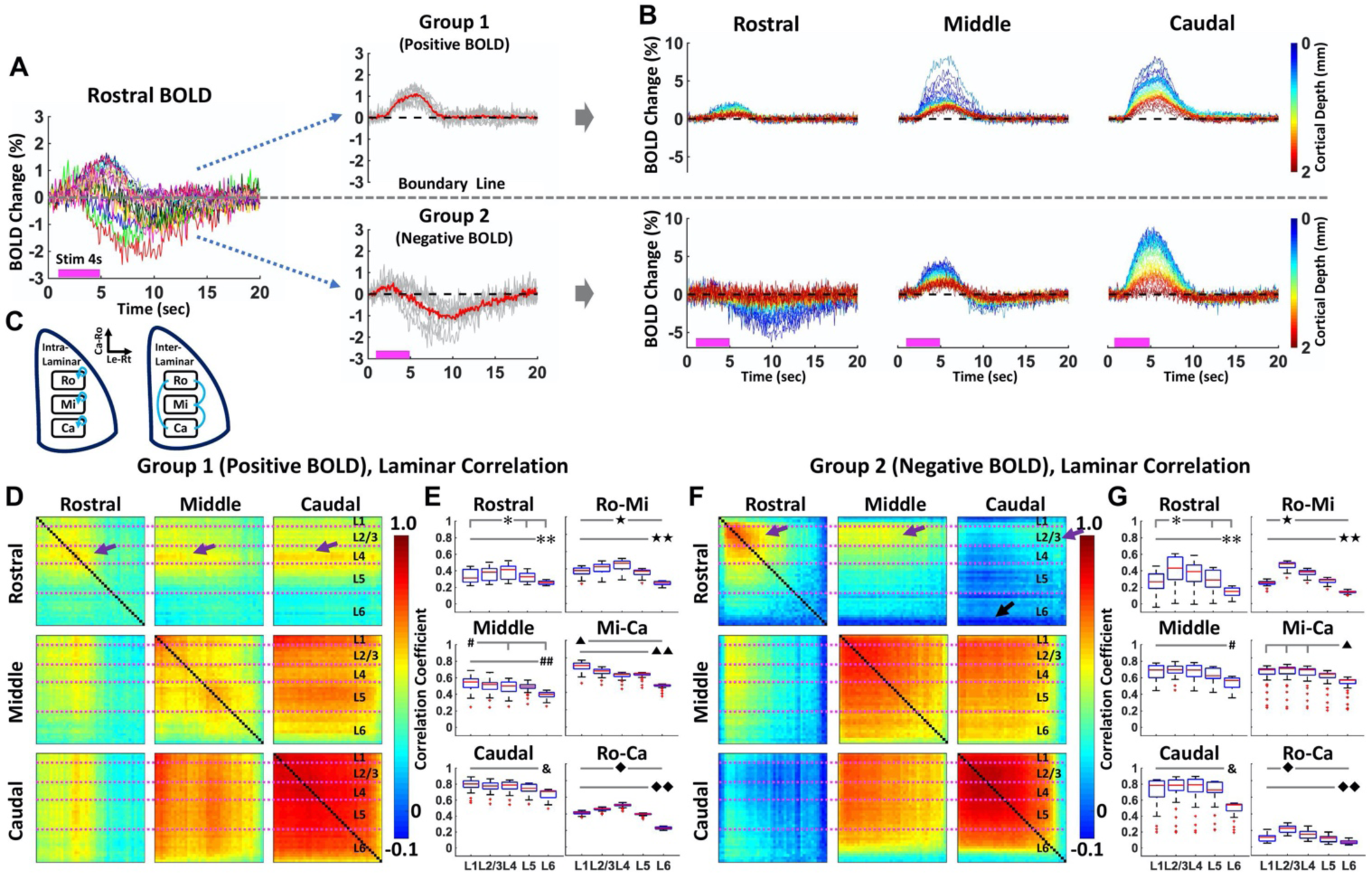
**A**. *Left*: Average BOLD changes (every 20 sec) of the average voxel time series (average of 40 voxels) showing both positive and negative BOLD responses in the rostral region with different trials (n = 18 trials of 4 rats). The individual colors represent individual trials. *Right*: Grouping based on either the positive or negative BOLD response in the rostral region. Group 1 (11 trials) has positive BOLD, and group 2 (7 trials) has negative BOLD. **B**. Three different average BOLD time series across the cortical depth (0 - 2 mm, 40 lines) in the rostral, middle, and caudal regions of group 1 and 2. **C**. Schematic illustration of intra- and inter-laminar correlation among the three cortical regions. **D-G**. Group-averaged results representing intra- and inter-laminar correlation maps in group 1 (**D** and **E**) and group 2 (**F** and **G**). In a 3 × 3 matrx (**D** and **F)**, diagonal and off-diagonal blocks represent intra- and inter-laminar correlation maps, respectively. L4 in group 1 and L2/3 in group 2 has higher correlation coefficients across the different cortical regions (purple arrows), showing significant difference (**E** and **G**) among all the layers. L6 shows significant difference in both groups while some parts of L6 in group 2 have negative correlation (**F**, black arrow). All statistic tests were performed with one-way ANOVA (post-hoc: p < 0.05, Bonferroni correction).

Next, the intra-and inter-laminar correlation maps (**Fig. 2C**) were calculated to obtain group-averaged results. In the 3 × 3 matrix (**Fig. 2D** and **2F**), the intra-laminar correlation maps were arranged on the diagonal blocks, while the inter-laminar correlation maps (i.e., rostral-middle, rostral-caudal, and middle-caudal correlation maps) were arranged on the off-diagonal blocks. From the intra-laminar correlation analysis, higher correlation coefficients were observed in superficial layers over deep layers of caudal and middle slices in both groups, but the rostral slice showed stronger correlation at Layer (L) 4 in group 1 versus L2/3 in group 2 (**Fig. 2E** and **2G**). Also, the inter-laminar correlation coefficients in group 1 showed higher values in L4 over other layers, presumably presenting robust BOLD signal correlation of FP-S1 through thalamocortical projections (25,41,42). In contrast, the correlation coefficients for Ro-Mi and Ro-Ca in group 2 showed higher values in L2/3, possibly presenting a negative BOLD signal-dominated correlation through corticocortical projection-mediated lateral inhibition (43-47). These results demonstrate distinct layer-specific correlation features underlying the positive and negative BOLD at the cortical areas adjacent to activated FP-S1, indicating brain state-dependent laminar fMRI responses across different trials.

### Mapping the laminar-specific functional connectivity in rs-fMRI

The MS-LS method could also be used to investigate the laminar-specific correlation features of rs-fMRI signals from the different cortial regions. **Fig. 3A** shows representative Z-score normalized time courses (average of 40 voxels) from the rostral, middle, and caudal slices, as well as the 2D line profile rs-fMRI maps as a function of time across the cortical depth, specifically presenting a highly synchronized slow oscillatory pattern. **Fig. 3B** shows the inter- and intra-laminar correlation maps of the rs-fMRI signals, presenting the highly correlated low-frequency signal fluctuation distributed in different cortical layers and across multipe cortical areas. In particular, the power spectral density (PSD) analysis of the rs-fMRI signal fluctuation shows peaked oscillatory powers at 0.01-0.02 Hz across different cortial areas (**Fig. 3C**), which is consistent with the previous study in anesthetized rats (48). Based on intra-laminar cross-correlation analysis, earlier lag times were detected in the middle layers (L2/3, 4, 5, median: −0.157, −0.207, −0.171 sec), whereas significantly later lag times were detected in the deep layer (L6, median: 0.216 sec) when compared to the mean BOLD signal of all the layers (**Fig. 3D**). To examine how the rs-fMRI signal propagated across different cortical areas, we calculated inter-laminar lag times by comparing layer-wise BOLD signals between slices of Ro-Mi, Mi-Ca, and Ro-Ca (**Fig. 3E**). The lag times were not uniform across different layers. For the Ro-Ca case, it shows clear fMRI signal propagation from the caudal to the rostral slice at L2/3 and 4 (−0.806 ± 0.703 sec, −0.577 ± 0.651 sec). Similarly, the Ro-Mi case shows earlier fMRI signal fluctuation at L2/3 to 6 from the middle to the rostral slice, but opposite at L1 (0.223 ± 0.810 sec), which may be caused by the large draining veins distributed across the middle slice (**Fig. 3A**). For the Mi-Ca case, the different lag time was only observed at L6, showing significantly later fMRI signal fluctuation at the caudal slice (0.777 ± 0.616 sec). These results demonstrate resting-state laminar-specific temporal correlation features across different cortical areas, which can be attributed to underlying either neuromodulation or vascular distribution across different cortical layers.

**Figure 3.**
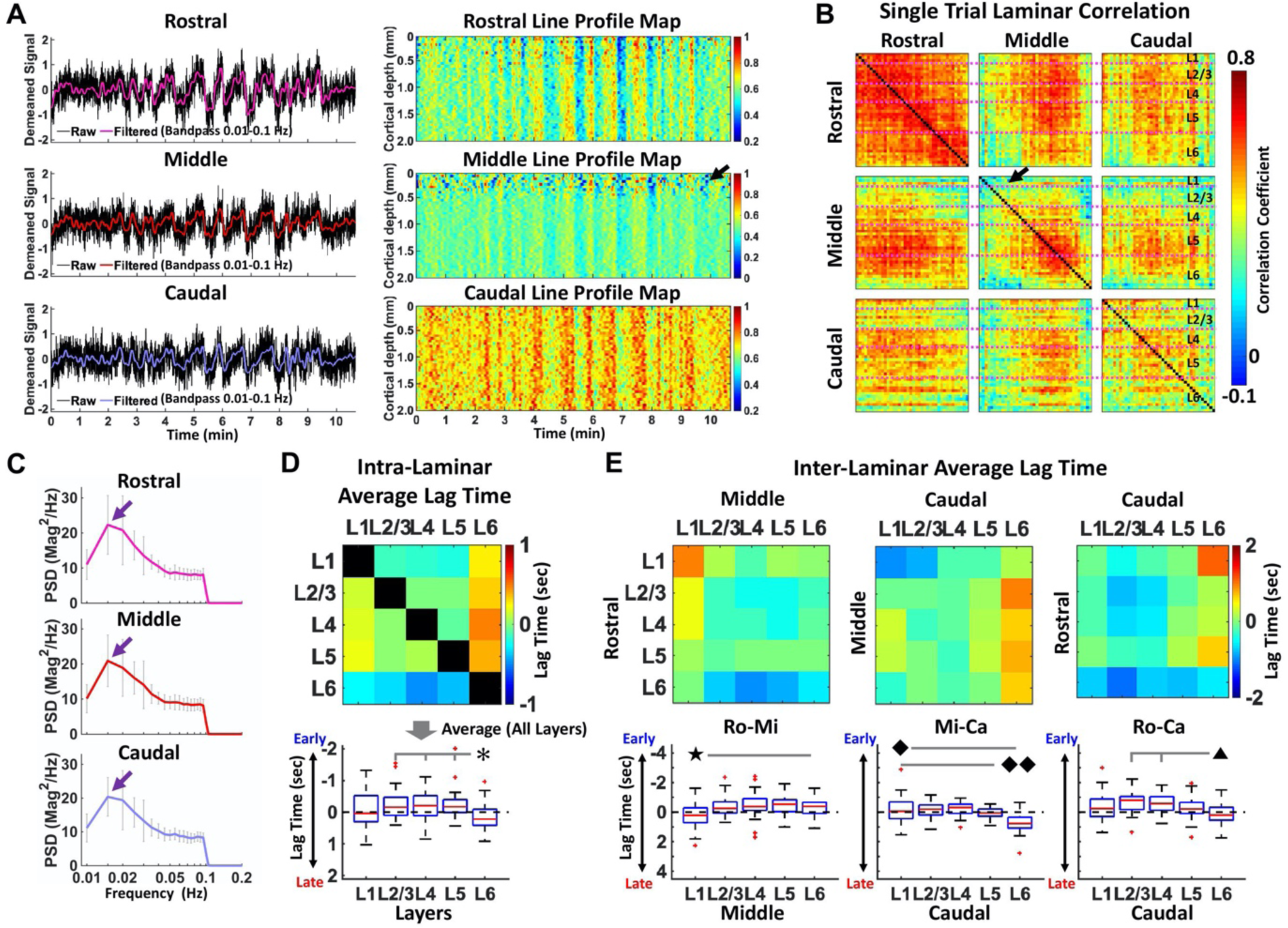
**A**. *Left*: Demeaned rs-fMRI time series of raw and filtered (bandpass: 0.01-0.1 Hz) data (average of 40 voxels) in rostral (upper), middle (middle), and caudal (lower) cortical regions from one representative trial (10 min 40 sec). *Right*: Normalized line profile maps showing the laminar-specific responses across the cortical depth (0 – 2 mm, 40 voxels) in the three different regions. The maps were processed by band-pass filtering (0.01-0.1 Hz). **B**. Laminar-specific correlation maps from the same trial in (**A**). In a 3 × 3 matrx, diagonal and off-diagonal blocks represent intra- and inter-laminar correlation maps, respectively. The black arrows (**A** and **B**) in the middle region indicates the BOLD effects of large draining vein around the superficial layers (L1 and L2/3) **C**. The PSDs of Z-score normalized fMRI time series (average of 40 voxels) showing the peaked low-frequency power at 0.01-0.02 Hz (purple arrows). **D** and **E**. Average results (n = 18 trials of 4 rats) representing layer-wise intra- (**D**) and inter-laminar (**E**) lag times. **D**. The middle layers (L2/3, 4, 5) show earlier lag times whereas L6 shows significantly later lag times. **E**. In Ro-Mi, Mi-Ca, and Ro-Ca lag times, L2/3, 4, 5 had ealier lag times (except L5 in Mi-Ca) whereas L1 and 6 had significantly later (except L1 in Mi-Ca) lag times, clearly showing the caudal to the rostral rs-fMRI signal propagation via L2/3 and L4. All significant tests were performed with one-way ANOVA (post-hoc: p-value <0.05, Bonferroni correction).

## DISCUSSION

In this work, we applied the MS-LS method to identify intra- and inter-laminar correlation patterns of evoked and resting-state fMRI signals with high spatial and temporal resolution across multiple cortical regions. Distinct laminar-specific correlation patterns in the cortex adjacent to the activated FP-S1 were detected based on either positive or negative BOLD responses, which may be regulated by altered brain states across different fMRI trials in anesthetized rats (36,49-51). The MS-LS mapping scheme is also applied to map the laminar correlation pattern of low frequency rs-fMRI signal fluctuation, presenting sub-second lag times of rs-fMRI signal propagation through cortical layers and across different cortical regions given the fast sampling rate.

An interesting observation of evoked laminar fMRI signal using the MS-LS method is the negative BOLD responses detected in the rostral region adjacent to FP-S1. Specifically, this temporal feature of the negative BOLD signal is different from the initial dip of positive BOLD as reported previously (52,53). It remains unclear whether the negative BOLD response has vascular or neuronal origins (54-64). As indicated by extensive studies, the BOLD signal relies on the integrated interactions of cerebral blood flow and volume (i.e., CBF and CBV) changes with metabolic rate of oxygen consumption (CMRO2) which are in principle caused by balanced proportional changes in both excitatory and inhibitory neuronal activity (65-67). Putative mechanisms of negative BOLD related to specific hemodynamic responses suggest two sources: i) the remaining elevation of CBV in contrast to returning to baseline CBF and CMRO2 (54-57), ii) the decoupling between CBF and CMRO2 during or after stimulation (57-59). Also, based on the vascular blood supply, a local blood-stealing effect from adjacent cortices to provide more blood to the most activated region is also proposed (60,61). With direct neuronal activity recording, the suppression of neuronal activity in the adjacent cortical regions due to lateral inhibition has also been reported (62-64). The first two mechanisms (i and ii) for signal decrease below baseline following the BOLD activation profile are likely explained by the prolonged post-stimulus undershoot in activated brain regions. As shown in **Fig. 2B**, while the negative BOLD mainly occured at the superficial layer where large arteries may contribute to CBV signal changes, robust negative BOLD signal was also detected at the L2/3 (blue color in the color bar). Also, the negative BOLD signal was maintained with longer spreading function than typical positive BOLD responses excluded the only contribution from CBV and CBF effects, suggesting a sustained corticocortical post-synaptic inhibition (**Fig. S1 A**) (47,68). It is also noteworthy that the negative BOLD group (group 2) shows the salient biphasic HRF of the positive BOLD, of which the post-stimulus undershoot may be also caused by the decreased neuronal activity (43).

Using the MS-LS method, we also detected the robust rs-fMRI slow oscillation pattern across cortical layers of three slices with a peak frequency power at 0.01-0.02 Hz. Previous studies with simultaneous fMRI and optical fiber-based Ca^2+^ recording in anesthetized rats have reported that neuronal calcium oscillations underlie the low frequency rs-fMRI signal fluctuation near 0.01-0.04 Hz (48). Also, intrinsic astrocytic Ca^2+^ transients have been reported to mediate global negative BOLD signals contributing to the low-frquency rs-fMRI signal fluctuation (69), suggesting brain state-dependent global neuromodulation of the ultra-slow oscillatory patterns (70-72). Moreover, negative global rs-fMRI during ultra-slow oscillation is also linked with pupil dynamics, showing converged effect of arousal state fluctuation and autonomus regulation (73-77). In contrast to the brain-wide rs-fMRI mapping of previous studies, we revealed laminar-specific signal propagation of the low frequency rs-fMRI signal fluctuation (**Fig. 3E**). Interestingly, the signal propagation direction varied at the different cortical layers. We observed more uniform lag times from L2/3 to L5 between different cortical regions, but largely varied lag times at L1 and L6. It is plausible that rs-fMRI signal oscillation in L2/3 to L5 is primarily driven by the global neuromodulation through subcortical projections (78), but the signal oscillation in L1 and L6 more likely relies on the different vascular density distribution that contributes to dynamic BOLD responses (79-82).

Technical limitations pertaining to MS-LS acquisition should be considered for future work. As a modification to the conventional line-scanning method (25), two saturation RF pulses were followed by three excitation RF pulses to acquire fMRI signals from three slices using MS-LS method. Given a certain set of sequence parameters (e.g., TE, readout bandwidth, etc), imperfect performances of the saturation RF pulses result in contaminating fMRI signals, due to aliasing artifacts that arise from fast T1 relaxation of tissue signals in outside of the FOV. It can be avoided by applying a region-of-interest selective refocusing RF pulse (83-85) instead of saturation RF pulses. Moreover, in the interleaved acquisition of MS-LS, we have longer TR (100 ms in this work compared to 50 ms TR of the conventional method), thus preserving readout bandwidth due to tradeoff between tSNR enhancement and minimum TR. For the future work, implantable inductive coils (86) or wireless amplified NMR detector (87,88) can be applied to enhance tSNR and to achieve a fast sampling rate simultaneously.

## METHODS

### Animal preparation

The study was performed in accordance with the German Animal Welfare Act (TierSchG) and Animal Welfare Laboratory Animal Ordinance (TierSchVersV), in full compliance with the guidelines of the EU Directive on the protection of animals used for scientific purposes (2010/63/EU). The study was reviewed by the ethics commission (§15 TierSchG) and approved by the state authority (Regierungspräsidium, Tübingen, Baden-Württemberg, Germany). A 12-12 hour on/off lighting cycle was maintained to assure undisturbed circadian rhythm. Food and water were obtainable ad libitum. A total of 4 male Sprague–Dawley rats were used in this study.

Anesthesia was first induced in the animal with 5% isoflurane in the chamber. The anesthetized rat was intubated using a tracheal tube. A mechanical ventilator (SAR-830, CWE, USA) was used to ventilate animals throughout the whole experiment. Femoral arterial and venous catheterization was performed with polyethylene tubing for blood sampling, drug administration, and constant blood pressure measurements. After the surgery, isoflurane was switched off, and a bolus of the anesthetic alpha-chloralose (80 mg/kg) was infused intravenously. After the animal was transferred to the MRI scanner, a mixture of alpha-chloralose (26.5 mg/kg/h) and pancuronium (2 mg/kg/h) was constantly infused to maintain the anesthesia level for reduced motion artifacts.

### EPI fMRI acquisition

All data sets from rats were acquired using a 14.1T/26 cm (Magnex, Oxford) horizontal bore magnet with an Avance III console (Bruker, Ettlingen) and a 12-cm diameter gradient system (100 G/cm, 150 µs rising time). A home-made transceiver surface coil with 6-mm diameter was used on the rat brain in all experiments. For the functional map of BOLD activation (**Fig. 1A)**, a 3D gradient-echo EPI sequence was acquired with the following parameters: TR/TE 1500/11.5 ms, FOV 1.92 × 1.92 × 1.92 cm^3^, matrix size 48 × 48 × 48, spatial resolution 0.4 × 0.4 × 0.4 mm^3^. A high order (*e*.*g*., 2^nd^ or 3^rd^ order) shimming was applied to reduce the main magnetic field (B0) inhomogeneities at the region-of-interest. For anatomical reference of the activated BOLD map, a RARE sequence was applied to acquire 48 coronal images with the same geometry as that of the EPI images. The fMRI design paradigm for each trial comprised of 200 dummy scans to reach steady-state, 10 pre-stimulation scans, 3 scans during stimulation, and 12 post-stimulation scans with a total of 8 epochs.

### MS-LS acquisition

GRE-based MS-LS datasets were acquired in anesthetized rats for evoked and rs-fMRI. The MS-LS method was applied by increasing the slice dimension (1 to 3) to record fMRI signals in three different cortical regions covering FP-S1 (i.e., rostral, middle, and caudal cortices) and using two saturation slices to avoid aliasing artifacts along the phase-encoding direction (**Fig. 1A**). The phase-encoding gradient was turned off to acquire line profiles (**Fig. 1C**). Laminar-specific fMRI responses from the three cortices were acquired along the frequency-encoding direction with 50-µm spatial resolution. The following acquisition parameters were used: TR/TE 100/9 ms, TA 10 min 40 sec, FA 50°, slice thickness 1.2 mm, slice gap 1.5 mm, FOV 6.4 × 3.2 mm^2^, and matrix 128 × 32. The fMRI design paradigm for each epoch consisted of 1 second pre-stimulation, 4 seconds stimulation, and 15 seconds post-stimulation for a total of 20 seconds. A total of 6400 lines (i.e., 10 min 40 sec) in each cortex were acquired in every single trial for evoked and rs-fMRI. Evoked BOLD activation was identified by performing electrical stimulation to the left forepaw (300 µs duration at 2.5 mA repeated at 3 Hz for 4 seconds).

### Data Analysis

All signal processing and analyses were implemented in MATLAB software (Mathworks, Natick, MA) and Analysis of Functional NeuroImages software (89) (AFNI, NIH, USA). For evoked fMRI analysis in **Fig. 1A**, the hemodynamic response function (HRF) used was the default of the block function of the linear program 3dDeconvolve in AFNI. BLOCK (L, 1) computes a convolution of a square wave of duration L and makes a peak amplitude of block response = 1, with *g*(*t*) = *t*^4^*e*^−*t*^/[4^4^*e*^−4^]. Each beta weight represents the peak height of the corresponding BLOCK curve for that class. The HRF model is defined as follows:

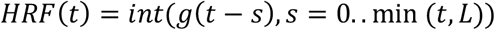

Cortical surfaces were determined based on signal intensities of fMRI line profiles (**Fig. 1C**). The detailed processing procedure was conducted as described in the previous line-scanning study (25). For **Fig. 1D** and **3A**, demeaned fMRI time courses were used as follows: (x - µ), where x was the original fMRI time courses and µ was the mean of the time courses. The line profile map concatenated with multiple line-scanning fMRI profiles was normalized by its maximum intensity. Average BOLD time series and percentage changes were defined as (S-S0)/S0 × 100 %, where S was the BOLD signal and S0 was the baseline. S0 was obtained by averaging the fluctuation signal in the 1-second pre-stimulation window in evoked fMRI that was repeated every 20 seconds with the whole time series (640 sec). The BOLD time series in each ROI were detrended and bandpass filtered (0.01-0.1 Hz, FIR filter) before analyzing line profile maps, correlation coefficients, PSDs, and cross-correlation.

Temporal correlation analysis which is generally accepted as an indicator of corticocortical functional interaction, *e*.*g*., increased correlation is thought to reflect increased functional connectivity between two brain sites (90,91). In this study, the correlation analysis was employed as an important indicator of laminar-specific functional connectivity across multiple cortical regions. Laminar-specific fMRI time courses were used and converted to the bandpass filtered time courses. The correlation is defined as follows:

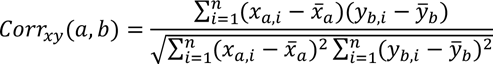

where *x, y* indicates filtered fMRI time series (n time points) and *a, b* indicates a voxel from one cortical region. The correlation was calculated by ‘corr’ function in MATLAB (Type: Pearson, Rows: pairwise).

For resting-state PSD analysis, fMRI time series was converted to the Z-score normalized time series. Subsequently, the converted time series was used to compare the frequency responses among the different regions avoiding the dependency of the difference in signal amplitudes (**Fig. 3C**). The Z-score normalized time courses were calculated as follows: (x - µ) / σ, where x was original fMRI time courses and µ, σ were the mean and the standard deviation of the time courses, respectively (zscore function in Matlab). PSDs were calculated by Welch’s estimation method (pwelch function in MATLAB, FFT length: 2000, overlap: 50%).

For lag time calculation in rs-fMRI, voxel-wise lag times were calculated by analyzing cross-correlation coefficients across all cortical layers or across different cortical regions (xcorr function in MATLAB). One laminar-specific lag time was determined as the time point with the maximal cross-correlation coefficient (**Fig. 3D** and **E**). The boundaries of different cortical layers were determined as provided in the previous line-scanning study (25).

For statistical analysis in evoked and rs-fMRI, one-way ANOVA was performed to compare laminar-specific values using the post-hoc test and to determine whether there were statistically significant differences among the associated population means in different cortical layers or areas. Student t-test was performed to divide all the trials into two groups (**Fig. S1 B)**. The p-values < 0.05 were considered statistically significant.

## Data availability

The data that support the findings of this study are available from the corresponding authors upon request.

## Code availability

The related sigal processing codes are available from the corresponding author upon reasonable request.

## Competing interests

The authors declare no competing interests.

## ACKNOWLEDGEMENTS

This research was supported by NIH Brain Initiative funding (RF1NS113278-01, R01 MH111438-01), and the S10 instrument grant (S10 MH124733) to Martinos Center, German Research Foundation (DFG) Yu215/3-1, BMBF 01GQ1702, and the internal funding from Max Planck Society. This project has received funding from the European Union Framework Programme for Research and Innovation Horizon 2020 (2014-2020) under the Marie Sklodowska-Curie Grant Agreement No.896245. We thank Dr. R. Pohmann, Dr. J. Engelmann, Dr. N. Avdievitch, and Ms. H. Schulz for technical support, Dr. P. Douay, Ms. R. König, and Ms. M. Pitscheider for animal support, the AFNI team for the software support.

## Supplementary information

**Figure S1.**
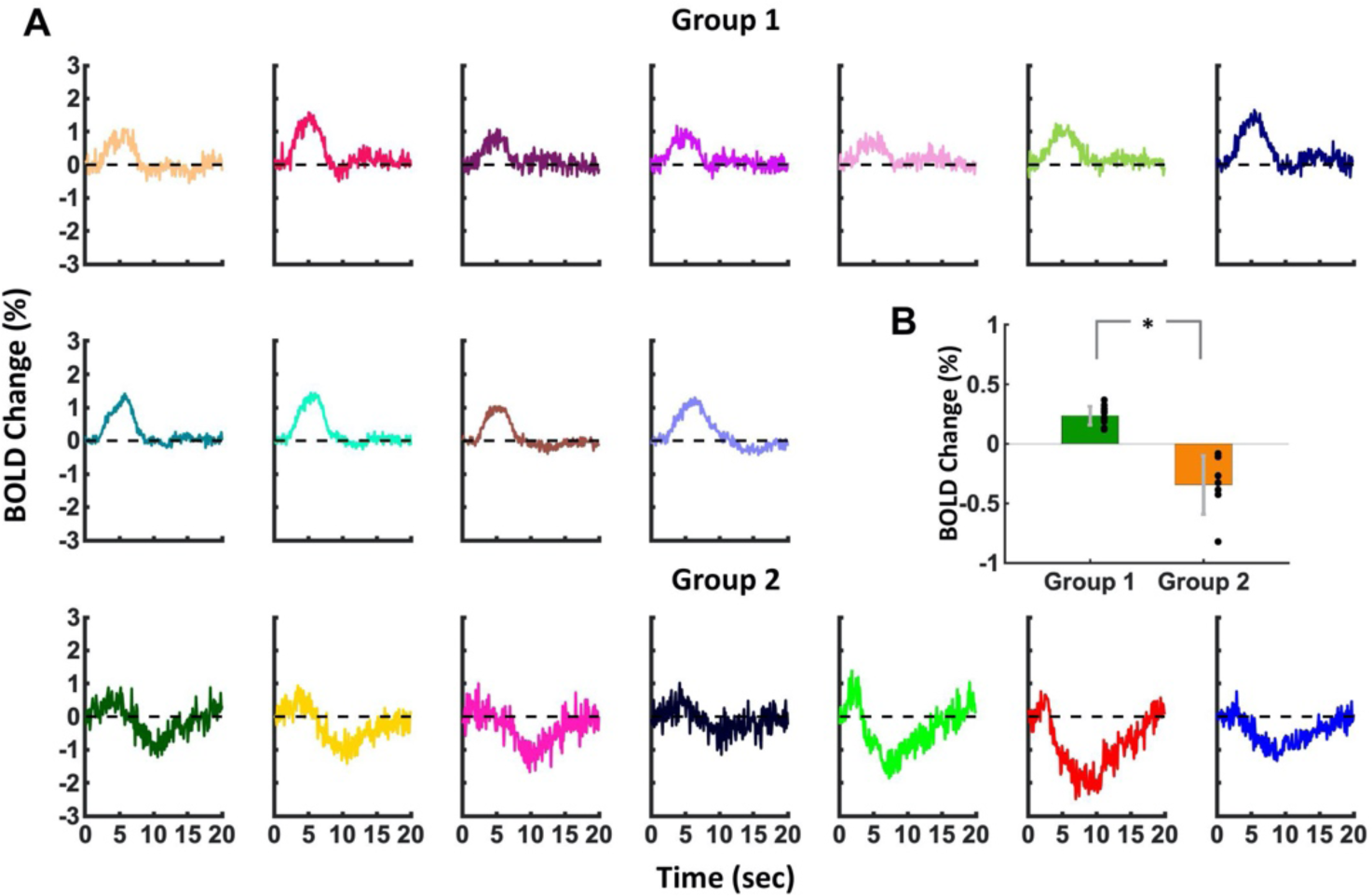
Evoked average BOLD time coures in the two groups. **A**. Average BOLD time courses (mean epoch with 20 sec) of average voxel (a total of 40 voxels) representing that group 1 (11 trials) has positive BOLD while group 2 (7 trials) has negative BOLD. **B**. Student t-test result with all the trials showing group 1 and 2 are significantly different (*p = 1.5055*10^-6).

## Notes

### Competing Interest Statement

The authors have declared no competing interest.

### Summary of Updates

Typo correction in Abstract

## REFERENCES

1. Biswal B, Yetkin FZ, Haughton VM, Hyde JS. Functional connectivity in the motor cortex of resting human brain using echo-planar MRI. Magn Reson Med 1995;34(4):537–541.

2. Biswal BB, Mennes M, Zuo XN, Gohel S, Kelly C, Smith SM, Beckmann CF, Adelstein JS, Buckner RL, Colcombe S, Dogonowski AM, Ernst M, Fair D, Hampson M, Hoptman MJ, Hyde JS, Kiviniemi VJ, Kotter R, Li SJ, Lin CP, Lowe MJ, Mackay C, Madden DJ, Madsen KH, Margulies DS, Mayberg HS, McMahon K, Monk CS, Mostofsky SH, Nagel BJ, Pekar JJ, Peltier SJ, Petersen SE, Riedl V, Rombouts SA, Rypma B, Schlaggar BL, Schmidt S, Seidler RD, Siegle GJ, Sorg C, Teng GJ, Veijola J, Villringer A, Walter M, Wang L, Weng XC, Whitfield-Gabrieli S, Williamson P, Windischberger C, Zang YF, Zhang HY, Castellanos FX, Milham MP. Toward discovery science of human brain function. Proc Natl Acad Sci U S A 2010;107(10):4734–4739.

3. Biswal BB. Resting state fMRI: a personal history. Neuroimage 2012;62(2):938–944.

4. Buckner RL, Krienen FM, Yeo BT. Opportunities and limitations of intrinsic functional connectivity MRI. Nat Neurosci 2013;16(7):832–837.

5. Fox MD, Raichle ME. Spontaneous fluctuations in brain activity observed with functional magnetic resonance imaging. Nat Rev Neurosci 2007;8(9):700–711.

6. Turner R, Le Bihan D, Moonen CT, Despres D, Frank J. Echo-planar time course MRI of cat brain oxygenation changes. Magn Reson Med 1991;22(1):159–166.

7. Smith SM, Fox PT, Miller KL, Glahn DC, Fox PM, Mackay CE, Filippini N, Watkins KE, Toro R, Laird AR, Beckmann CF. Correspondence of the brain’s functional architecture during activation and rest. P Natl Acad Sci USA 2009;106(31):13040–13045.

8. Yeo BTT, Krienen FM, Sepulcre J, Sabuncu MR, Lashkari D, Hollinshead M, Roffman JL, Smoller JW, Zoller L, Polimeni JR, Fischl B, Liu HS, Buckner RL. The organization of the human cerebral cortex estimated by intrinsic functional connectivity. Journal of Neurophysiology 2011;106(3):1125–1165.

9. Beckmann CF, DeLuca M, Devlin JT, Smith SM. Investigations into resting-state connectivity using independent component analysis. Philos T R Soc B 2005;360(1457):1001–1013.

10. Preibisch C, Castrillon JG, Buhrer M, Riedl V. Evaluation of Multiband EPI Acquisitions for Resting State fMRI. Plos One 2015;10(9).

11. Smith SM, Vidaurre D, Beckmann CF, Glasser MF, Jenkinson M, Miller KL, Nichols TE, Robinson EC, Salimi-Khorshidi G, Woolrich MW, Barch DM, Ugurbil K, Van Essen DC. Functional connectomics from resting-state fMRI. Trends in Cognitive Sciences 2013;17(12):666–682.

12. Huber L, Handwerker DA, Jangraw DC, Chen G, Hall A, Stuber C, Gonzalez-Castillo J, Ivanov D, Marrett S, Guidi M, Goense J, Poser BA, Bandettini PA. High-Resolution CBV-fMRI Allows Mapping of Laminar Activity and Connectivity of Cortical Input and Output in Human M1. Neuron 2017;96(6):1253-+.

13. Sharoh D, van Mourik T, Bains LJ, Segaert K, Weber K, Hagoort P, Norris DG. Laminar specific fMRI reveals directed interactions in distributed networks during language processing. P Natl Acad Sci USA 2019;116(42):21185–21190.

14. Finn ES, Huber L, Jangraw DC, Molfese PJ, Bandettini PA. Layer-dependent activity in human prefrontal cortex during working memory. Nature Neuroscience 2019;22(10):1687-+.

15. Kashyap S, Ivanov D, Havlicek M, Sengupta S, Poser BA, Uludag K. Resolving laminar activation in human V1 using ultra-high spatial resolution fMRI at 7T. Sci Rep-Uk 2018;8.

16. Yu Y, Huber L, Yang J, Jangraw DC, Handwerker DA, Molfese PJ, Chen G, Ejima Y, Wu J, Bandettini PA. Layer-specific activation of sensory input and predictive feedback in the human primary somatosensory cortex. Sci Adv 2019;5(5):eaav9053.

17. Goense JBM, Logothetis NK. Laminar specificity in monkey V1 using high-resolution SE-fMRI. Magn Reson Imaging 2006;24(4):381–392.

18. Silva AC, Koretsky AP. Laminar specificity of functional MRI onset times during somatosensory stimulation in rat. Proceedings of the National Academy of Sciences of the United States of America 2002;99(23):15182–15187.

19. Lewis LD, Setsompop K, Rosen BR, Polimeni JR. Fast fMRI can detect oscillatory neural activity in humans. P Natl Acad Sci USA 2016;113(43):E6679–E6685.

20. Agrawal U, Brown EN, Lewis LD. Model-based physiological noise removal in fast fMRI. Neuroimage 2020;205.

21. Gil R, Fernandes FF, Shemesh N. Neuroplasticity-driven timing modulations revealed by ultrafast functional magnetic resonance imaging. Neuroimage 2021;225.

22. Pais-Roldan P, Biswal B, Scheffler K, Yu X. Identifying Respiration-Related Aliasing Artifacts in the Rodent Resting-State fMRI. Front Neurosci 2018;12:788.

23. Birn RM, Diamond JB, Smith MA, Bandettini PA. Separating respiratory-variation-related fluctuations from neuronal-activity-related fluctuations in fMRI. Neuroimage 2006;31(4):1536–1548.

24. Murphy K, Birn RM, Bandettini PA. Resting-state fMRI confounds and cleanup. Neuroimage 2013;80:349–359.

25. Yu X, Qian C, Chen DY, Dodd SJ, Koretsky AP. Deciphering laminar-specific neural inputs with line-scanning fMRI. Nat Methods 2014;11(1):55–58.

26. Albers F, Schmid F, Wachsmuth L, Faber C. Line scanning fMRI reveals earlier onset of optogenetically evoked BOLD response in rat somatosensory cortex as compared to sensory stimulation. Neuroimage 2018;164:144–154.

27. Nunes D, Gil R, Shemesh N. A rapid-onset diffusion functional MRI signal reflects neuromorphological coupling dynamics. Neuroimage 2021;231:117862.

28. Raimondo L, Knapen T, Oliveira I, Yu X, van der Zwaag W, Siero J. Preliminary results of functional line-scanning in humans: submillimeter, subsecond resolution evoked responses. 36th Annual Scientific Meeting of the European Society for Magnetic Resonance in Medicine and Biology 2019;S22.06:S334.

29. Siero J, Oliveira I, Choi S, Yu X. Implementing human line-scanning fMRI: Initial results of ultra-high temporal and spatial resolution fMRI. Proc Intl Soc Mag Reson Med 2019;27:3933.

30. Morgan AT, Nothnagel N, Petro LS, Goense J, Muckli L. High-resolution line-scanning reveals distinct visual response properties across human cortical layers. bioRxiv 2020:2020.2006.2030.179762.

31. Balasubramanian M, Mulkern RV, Neil JJ, Maier SE, Polimeni JR. Probing in vivo cortical myeloarchitecture in humans via line-scan diffusion acquisitions at 7 T with 250-500 micron radial resolution. Magnetic resonance in medicine: official journal of the Society of Magnetic Resonance in Medicine / Society of Magnetic Resonance in Medicine 2021;85(1):390–403.

32. Jung WB, Shim HJ, Kim SG. Mouse BOLD fMRI at ultrahigh field detects somatosensory networks including thalamic nuclei. Neuroimage 2019;195:203–214.

33. Kim T, Masamoto K, Fukuda M, Vazquez A, Kim SG. Frequency-dependent neural activity, CBF, and BOLD fMRI to somatosensory stimuli in isoflurane-anesthetized rats. Neuroimage 2010;52(1):224–233.

34. Nair G, Duong TQ. Echo-planar BOLD fMRI of mice on a narrow-bore 9.4 T magnet. Magnet Reson Med 2004;52(2):430–434.

35. Kim T, Masamoto K, Fukuda M, Vazquez A, Kim SG. Frequency-dependent neural activity, CBF, and BOLD fMRI to somatosensory stimuli in isoflurane-anesthetized rats. Neuroimage 2010;52(1):224–233.

36. Devor A, Tian P, Nishimura N, Teng IC, Hillman EM, Narayanan SN, Ulbert I, Boas DA, Kleinfeld D, Dale AM. Suppressed neuronal activity and concurrent arteriolar vasoconstriction may explain negative blood oxygenation level-dependent signal. J Neurosci 2007;27(16):4452–4459.

37. Boorman L, Kennerley AJ, Johnston D, Jones M, Zheng Y, Redgrave P, Berwick J. Negative blood oxygen level dependence in the rat: a model for investigating the role of suppression in neurovascular coupling. J Neurosci 2010;30(12):4285–4294.

38. Goense J, Merkle H, Logothetis NK. High-resolution fMRI reveals laminar differences in neurovascular coupling between positive and negative BOLD responses. Neuron 2012;76(3):629–639.

39. de la Rosa-Rivera N, Ress D, Taylor AJ, Kim JH. Retinotopic variations of the negative blood oxygen-level dependent hemodynamic response function in human primary visual cortex. J Neurophysiol 2021.

40. Hu X, Le TH, Ugurbil K. Evaluation of the early response in fMRI in individual subjects using short stimulus duration. Magn Reson Med 1997;37(6):877–884.

41. Bopp R, Holler-Rickauer S, Martin KAC, Schuhknecht GFP. An Ultrastructural Study of the Thalamic Input to Layer 4 of Primary Motor and Primary Somatosensory Cortex in the Mouse. Journal of Neuroscience 2017;37(9):2435–2448.

42. El-Boustani S, Sermet BS, Foustoukos G, Oram TB, Yizhar O, Petersen CCH. Anatomically and functionally distinct thalamocortical inputs to primary and secondary mouse whisker somatosensory cortices. Nat Commun 2020;11(1).

43. Devor A, Tian PF, Nishimura N, Teng IC, Hillman EMC, Narayanan SN, Ulbert I, Boas DA, Kleinfeld D, Dale AM. Suppressed neuronal activity and concurrent arteriolar vasoconstriction may explain negative blood oxygenation level-dependent signal. Journal of Neuroscience 2007;27(16):4452–4459.

44. Kunori N, Takashima I. High-order motor cortex in rats receives somatosensory inputs from the primary motor cortex via cortico-cortical pathways. Eur J Neurosci 2016;44(11):2925–2934.

45. Thomson AM, Lamy C. Functional maps of neocortical local circuitry. Front Neurosci 2007;1(1):19–42.

46. Butovas S, Schwarz C. Spatiotemporal effects of microstimulation in rat neocortex: A parametric study using multielectrode recordings. Journal of Neurophysiology 2003;90(5):3024–3039.

47. Butovas S, Hormuzdi SG, Monyer H, Schwarz C. Effects of electrically coupled inhibitory networks on local neuronal responses to intracortical microstimulation. Journal of Neurophysiology 2006;96(3):1227–1236.

48. He Y, Wang MS, Chen XM, Pohmann R, Polimeni JR, Scheffler K, Rosen BR, Kleinfeld D, Yu X. Ultra-Slow Single-Vessel BOLD and CBV-Based fMRI Spatiotemporal Dynamics and Their Correlation with Neuronal Intracellular Calcium Signals. Neuron 2018;97(4):925-+.

49. Wang M, He Y, Sejnowski TJ, Yu X. Brain-state dependent astrocytic Ca(2+) signals are coupled to both positive and negative BOLD-fMRI signals. Proc Natl Acad Sci U S A 2018;115(7):E1647–E1656.

50. Parker DB, Razlighi QR. Task-evoked Negative BOLD Response and Functional Connectivity in the Default Mode Network are Representative of Two Overlapping but Separate Neurophysiological Processes. Sci Rep 2019;9(1):14473.

51. Kobayashi E, Bagshaw AP, Grova C, Dubeau F, Gotman J. Negative BOLD responses to epileptic spikes. Hum Brain Mapp 2006;27(6):488–497.

52. Tian PF, Teng IC, May LD, Kurz R, Lu K, Scadeng M, Hillman EMC, De Crespigny AJ, D’Arceuil HE, Mandeville JB, Marota JJA, Rosen BR, Liu TT, Boas DA, Buxton RB, Dale AM, Devor A. Cortical depth-specific microvascular dilation underlies laminar differences in blood oxygenation level-dependent functional MRI signal. P Natl Acad Sci USA 2010;107(34):15246–15251.

53. Siero JCW, Hendrikse J, Hoogduin H, Petridou N, Luijten P, Donahue MJ. Cortical depth dependence of the BOLD initial dip and poststimulus undershoot in human visual cortex at 7 Tesla. Magnet Reson Med 2015;73(6):2283–2295.

54. Harel N, Lee SP, Nagaoka T, Kim DS, Kim SG. Origin of negative blood oxygenation level-dependent fMRI signals. J Cerebr Blood F Met 2002;22(8):908–917.

55. Shmuel A, Yacoub E, Pfeuffer J, Van de Moortele PF, Adriany G, Hu XP, Ugurbil K. Sustained negative BOLD, blood flow and oxygen consumption response and its coupling to the positive response in the human brain. Neuron 2002;36(6):1195–1210.

56. Moraschi M, DiNuzzo M, Giove F. On the origin of sustained negative BOLD response. Journal of Neurophysiology 2012;108(9):2339–2342.

57. Yacoub E, Ugurbil K, Harel N. The spatial dependence of the poststimulus undershoot as revealed by high-resolution BOLD-and CBV-weighted fMRI. J Cerebr Blood F Met 2006;26(5):634–644.

58. Valabregue R, Aubert A, Burger J, Bittoun T, Costalat R. Relation between cerebral blood flow and metabolism explained by a model of oxygen exchange. J Cerebr Blood F Met 2003;23(5):536–545.

59. Buxton RB. Interpreting oxygenation-based neuroimaging signals: the importance and the challenge of understanding brain oxygen metabolism. Front Neuroenergetics 2010;2:8.

60. Smith AT, Williams AL, Singh KD. Negative BOLD in the visual cortex: Evidence against blood stealing. Human Brain Mapping 2004;21(4):213–220.

61. Bressler D, Spotswood N, Whitney D. Negative BOLD fMRI Response in the Visual Cortex Carries Precise Stimulus-Specific Information. Plos One 2007;2(5).

62. Shmuel A, Augath M, Oeltermann A, Logothetis NK. Negative functional MRI response correlates with decreases in neuronal activity in monkey visual area V1. Nature Neuroscience 2006;9(4):569–577.

63. Mullinger KJ, Mayhew SD, Bagshaw AP, Bowtell R, Francis ST. Evidence that the negative BOLD response is neuronal in origin: A simultaneous EEG-BOLD-CBF study in humans. Neuroimage 2014;94:263–274.

64. Pasley BN, Inglis BA, Freeman RD. Analysis of oxygen metabolism implies a neural origin for the negative BOLD response in human visual cortex. Neuroimage 2007;36(2):269–276.

65. Davis TL, Kwong KK, Weisskoff RM, Rosen BR. Calibrated functional MRI: Mapping the dynamics of oxidative metabolism. P Natl Acad Sci USA 1998;95(4):1834–1839.

66. Buxton RB, Uludag K, Dubowitz DJ, Liu TT. Modeling the hemodynamic response to brain activation. Neuroimage 2004;23:S220–S233.

67. Logothetis NK. What we can do and what we cannot do with fMRI. Nature 2008;453(7197):869–878.

68. Chen Y, Sobczak F, Pais-Roldan P, Schwarz C, Koretsky AP, Yu X. Mapping the Brain-Wide Network Effects by Optogenetic Activation of the Corpus Callosum. Cereb Cortex 2020;30(11):5885–5898.

69. Wang MS, He Y, Sejnowski TJ, Yu X. Brain-state dependent astrocytic Ca2+ signals are coupled to both positive and negative BOLD-fMRI signals. P Natl Acad Sci USA 2018;115(7):E1647–E1656.

70. Attwell D, Buchan AM, Charpak S, Lauritzen M, MacVicar BA, Newman EA. Glial and neuronal control of brain blood flow. Nature 2010;468(7321):232–243.

71. Petzold GC, Murthy VN. Role of Astrocytes in Neurovascular Coupling. Neuron 2011;71(5):782–797.

72. Poskanzer KE, Yuste R. Astrocytes regulate cortical state switching in vivo. P Natl Acad Sci USA 2016;113(19):E2675–E2684.

73. Chang C, Leopold DA, Scholvinck ML, Mandelkow H, Picchioni D, Liu X, Ye FQ, Turchi JN, Duyn JH. Tracking brain arousal fluctuations with fMRI. P Natl Acad Sci USA 2016;113(16):4518–4523.

74. Watson BO. Cognitive and Physiologic Impacts of the Infraslow Oscillation. Frontiers in Systems Neuroscience 2018;12.

75. Bradley MM, Miccoli L, Escrig MA, Lang PJ. The pupil as a measure of emotional arousal and autonomic activation. Psychophysiology 2008;45(4):602–607.

76. Liu X, de Zwart JA, Scholvinck ML, Chang CT, Ye FQ, Leopold DA, Duyn JH. Subcortical evidence for a contribution of arousal to fMRI studies of brain activity. Nat Commun 2018;9.

77. Ozbay PS, Chang C, Picchioni D, Mandelkow H, Chappel-Farley MG, van Gelderen P, de Zwart JA, Duyn J. Sympathetic activity contributes to the fMRI signal. Communications Biology 2019;2.

78. Constantinople CM, Bruno RM. Deep Cortical Layers Are Activated Directly by Thalamus. Science 2013;340(6140):1591–1594.

79. Koopmans PJ, Barth M, Norris DG. Layer-specific BOLD activation in human V1. Hum Brain Mapp 2010;31(9):1297–1304.

80. Markuerkiaga I, Barth M, Norris DG. A cortical vascular model for examining the specificity of the laminar BOLD signal. Neuroimage 2016;132:491–498.

81. Goense J, Bohraus Y, Logothetis NK. fMRI at High Spatial Resolution: Implications for BOLD-Models. Front Comput Neurosc 2016;10.

82. Schmid F, Barrett MJP, Jenny P, Weber B. Vascular density and distribution in neocortex. Neuroimage 2019;197:792–805.

83. Mansfield P, Maudsley AA, Baines T. Fast Scan Proton Density Imaging by Nmr. J Phys E Sci Instrum 1976;9(4):271–278.

84. Finsterbusch J, Frahm J. Gradient-echo line scan imaging using 2D-Selective RF excitation. J Magn Reson 2000;147(1):17–25.

85. Choi S, Zeng H, Pohmann R, Scheffler K, Yu X. Novel alpha-180 SE based LINE-scanning method (SELINE) for laminar-specific fMRI. Proc Intl Soc Mag Reson Med 2018;27:1166.

86. Inigo-Marco I, Isturiz J, Fernandez M, Nicolas MJ, Dominguez P, Bastarrika G, Valencia M, Fernandez-Seara MA. Imaging of Stroke in Rodents Using a Clinical Scanner and Inductively Coupled Specially Designed Receiver Coils. Annals of Biomedical Engineering 2020.

87. Qian CQ, Yu X, Chen DY, Dodd S, Bouraoud N, Pothayee N, Chen Y, Beeman S, Bennett K, Murphy-Boesch J, Koretsky A. Wireless Amplified Nuclear MR Detector (WAND) for High-Spatial-Resolution MR Imaging of Internal Organs: Preclinical Demonstration in a Rodent Model. Radiology 2013;268(1):228–236.

88. Qian W, Yu X, Qian CQ. Wireless Reconfigurable RF Detector Array for Focal and Multiregional Signal Enhancement. Ieee Access 2020;8:136594–136604.

89. Cox RW. AFNI: software for analysis and visualization of functional magnetic resonance neuroimages. Comput Biomed Res 1996;29(3):162–173.

90. Bandettini PA, Jesmanowicz A, Wong EC, Hyde JS. Processing strategies for time-course data sets in functional MRI of the human brain. Magn Reson Med 1993;30(2):161–173.

91. Friston KJ. Functional and effective connectivity: a review. Brain Connect 2011;1(1):13–36.

